# Stabilising selection and ecological trade-offs underpin coexistence in a tropical flora

**DOI:** 10.64898/2025.12.31.697164

**Authors:** Matti A. Niissalo, Jun Ying Lim, Jia Jun Ngiam, Sitaram Rajaraman, Le Min Choo, Joan Jing Yi Jong, Kang Min Ngo, Felicia Wei Shan Leong, Shawn Kaihekulani Yamauchi Lum, Henk J. Beentje, Sven Buerki, Martin W. Callmander, Junhao Chen, Matthias Seng En Chua, Rogier P.J. De Kok, Brigitta E.E. De Wilde-Duyfjes, Helena Duistermaat, Hans-Joachim Esser, Steven Fleck, S.K. Ganesan, Elliot M. Gardner, Mark Hughes, Sin Lan Koh, Paul K.F. Leong, Jana Leong-Škorničková, Stuart Lindsay, Yee Wen Low, Hock Keong Lua, Louise Neo, Caroline M. Pannell, Carmen Puglisi, Michele L. Rodda, Ivan A. Savinov, Wei Wei Seah, David A. Simpson, Joeri S. Strijk, Daniel C. Thomas, Ian M. Turner, Peter C. van Welzen, Peter Wilkie, Khoon Meng Wong, Chow Khoon Yeo, Victor A. Albert, Charles H. Cannon, Stuart J. Davies, Martin Lascoux, Charlotte Lindqvist, Puay Yok Tan, Timothy Utteridge, David A. Wardle, Kenneth B.H. Er, Peter R. Preiser, David J. Middleton, Gillian S. Khew, Jarkko Salojärvi

## Abstract

Tropical forests harbour the majority of global plant biodiversity^1,2^, yet the genomic mechanisms governing the assembly and maintenance of these communities remain poorly understood. Here, we assembled draft genomes for 499 angiosperm species from a lowland rainforest in Singapore, representing 67% of its flora, and integrated these with plant traits and comprehensive forest census data.

Across the community, most gene families evolve under stabilising selection, with copy numbers maintained near long-term optima that differ among ecological strategies. These niche-associated genomic attractor states provide a mechanism for convergent adaptation and species coexistence. Modelling stabilising selection also identified a strong trade-off between defence and growth, indicating that pathogen pressure constrains developmental diversification. Consistent with this, species-specific genome space was enriched for resistance genes and transposable elements.

In contrast, genomic processes structuring present-day plant community composition differ from those driving deep-time convergence. Genomic comparisons across forest types revealed stronger selection on defence-related pathways in old-growth primary forests and on growth-related processes in regenerating secondary forests, while community-level genomic profiles showed expansions in gene families associated with rapid responses to environmental fluctuations.

Stabilising selection therefore links population-level adaptation^3,4^ with long-term species diversification in the tropics. Niche similarity promotes long-term coexistence, whereas local community structure is shaped by more rapid ecological filtering driven by environmental change. Taken together, these two distinct evolutionary layers provide a genomic framework for understanding how hyperdiverse rainforest floras arise and persist.

## Introduction

The tropics harbour roughly two-thirds of global biodiversity^3^, and Southeast Asia (SEA) alone contains approximately 50,000 recorded vascular plant species – around 15% of Earth’s plant diversity within only ∼3% of its land area^3,4^. This exceptional richness reflects the region’s complex geological and climatic history^5^, including the gradual assembly of Laurasian, Indian and Sahul floras over the past 50 million years^6^. Although these biogeographic processes enabled large-scale dispersal and lineage accumulation, they do not by themselves explain the long-term coexistence of hyperdiverse communities nor reveal the evolutionary forces that have shaped tropical plant diversity through deep time.

An ecological perspective, by contrast, focuses on community assembly over much shorter intervals—typically no more than a few hundred years. During the initial ecological filtering stage, environmental conditions, resource distributions and biotic interactions restrict the subset of species able to recruit and coexist. Explanations for the resulting tropical community structure range from neutral, in which all species are functionally equivalent^5^, to niche-based models involving resource partitioning^7^, together with negative density-dependent effects^8^ exerted by herbivores and plant pathogens. Recent theoretical^9^ and simulation work^10^ further indicates that coexistence may arise even among species with highly similar resource-use profiles, generating clusters of functionally similar taxa (“species lumps”) under fluctuating resource regimes.

Such short-term filtering processes are expected to scale up over evolutionary timescales by shaping which lineages establish, persist and diversify. While large population sizes should in principle enhance the efficiency of selection in widespread species, thereby promoting ecological dominance, tropical forests also exhibit features such as high productivity^11,12^, intense biotic interactions, and, in angiosperms, tropical origins coupled with niche conservatism^11,13,14^ that could accelerate lineage diversification.

Despite their conceptual importance, the genomic foundations of these processes remain poorly resolved. Most genomic studies in tropical plants have focused on individual genera, providing limited insight into genome-wide evolutionary patterns at the community scale. Here we assembled and analysed draft genomes from 547 rainforest angiosperm species in a lowland dipterocarp forest in Southeast Asia, and combined the data with species traits and comprehensive forest census information. We show that genome evolution across this community is dominated by stabilizing selection, with ecological traits imposing distinct genomic optima. We further identify a pervasive defence–growth trade-off that limits the ability of species to track these optima, and find that species-specific genomic space is enriched for rapidly evolving resistance genes and transposable elements—implicating these mechanisms in recent diversification. Community assembly, in contrast, was distinct from the processes driven by trait-based convergence and shaped by functions linked with rapid environmental changes.

Together, these findings reveal two distinct evolutionary layers: long-term trait-driven genomic convergence towards niche-specific attractor states, and short-term ecological sorting driven by environmental fluctuations. This genomic flexibility offers a framework for understanding how hyperdiverse tropical floras are assembled and maintained.

### Sample collection and genome assembly – Cretacean origins of diversification

Singapore possesses one of the most comprehensively documented tropical floras, reflecting nearly two centuries of botanical research^15^. We focused on Bukit Timah Nature Reserve (BTNR; **Fig 1**), a small (164 hectares) but floristically rich lowland rainforest embedded within a highly urban landscape. The reserve comprises both old-growth primary forest that has never been clear-felled, and regenerating secondary forest established on previously disturbed land^16^. BTNR is among the most intensively studied forests in the region, with standardised and comprehensive ecological monitoring extending over three decades^17–20^.

**Figure 1:**
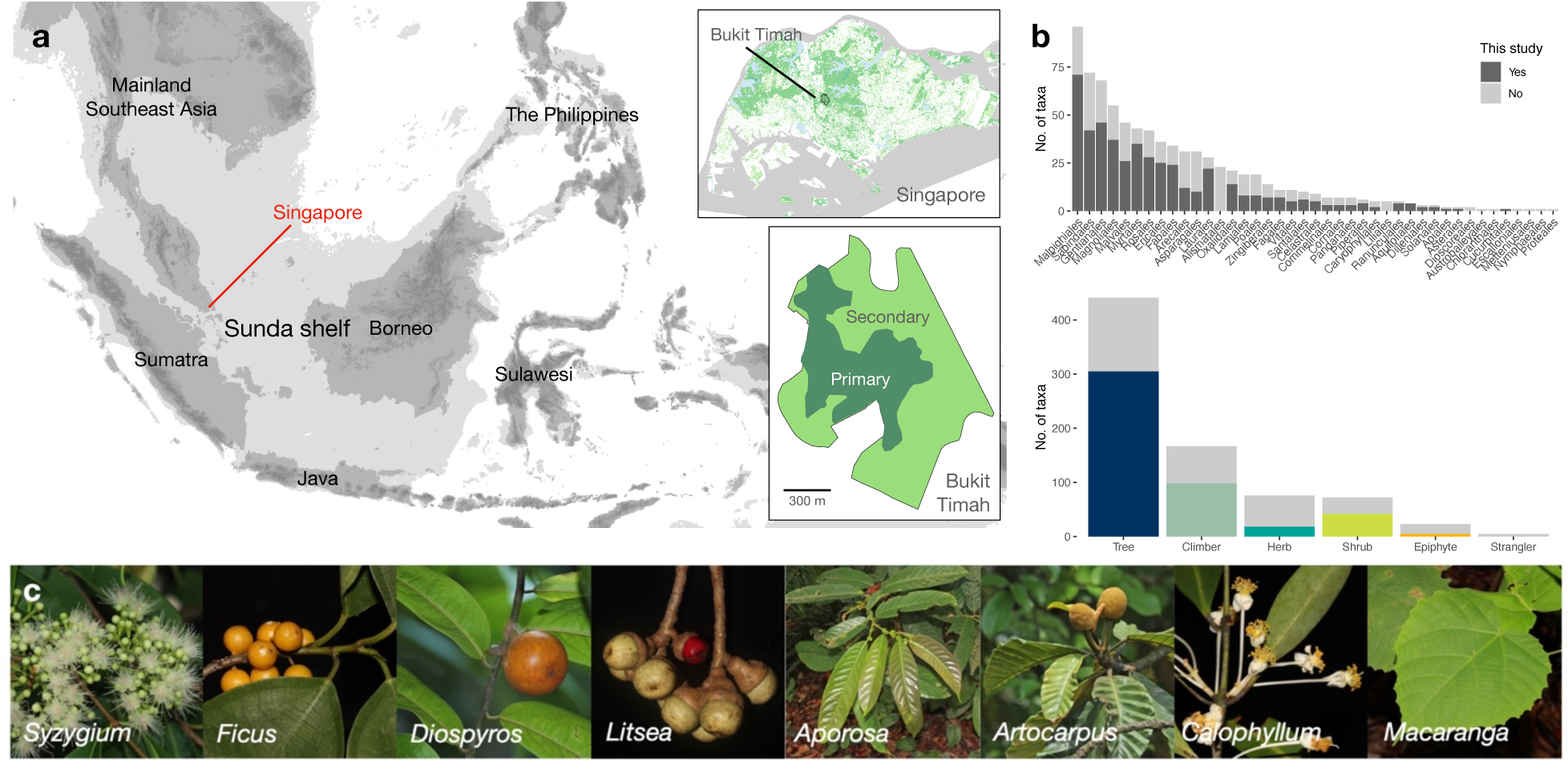
Sequencing the tropical forest of Bukit Timah Nature Reserve (BTNR). **a)** BTNR is situated in Singapore in the middle of the Sunda shelf, where plant populations experienced changes in connectivity with those of neighbouring land masses due to sea-level alterations during Pleistocene glacial-interglacial cycles (light grey areas extending beyond modern day coastlines represent areas that would have been mostly above sea level during glacial maxima, when sea level was 130 m below current levels). Like many forests in the region, BTNR has experienced past land-use changes, with more than half of its total area presently consisting of secondary forest (less mature, regrowth forest, characterised by less complex forest structure and fast growing, short-lived pioneer species). **b)** In terms of the relative coverage of species in our study, the angiosperm ordinal and life-form diversities are high. The dark grey bars in the top bar plot illustrate the species included in this study, while light grey demonstrates the total diversity encountered in BTNR. The bottom bar plot illustrates the life-form diversity, with the coloured fill demonstrating the species included in this study. **c)** The sampling includes some of the most species-rich genera in Southeast Asia. Here the eight largest genera in BTNR are illustrated, in descending order of number of species. Credit: Louise Neo (*Syzygium*, *Ficus*, *Aporosa*, *Artocarpus*, *Macaranga*) and Xin Yi Ng (*Diospyros*, *Litsea*, *Calophyllum*).

Leveraging data from these existing surveys and census efforts, we collected, identified, vouchered, sequenced and assembled the genomes of 547 native angiosperm species (**Supplementary Section 1;Table S1)**. Of these, 499 assemblies passed our quality control criteria (**Supplementary Section 2**), representing ∼60% of the entire angiosperm flora in the forest, and 79.4% of all large native trees (≥ 30 cm diameter at breast height). After contaminant removal, the assembly sizes ranged from 168 Mbp to 2.2 Gbp, with a median of 677 Mbp and median single-copy gene (BUSCO^21^) score of 87% (**Table S2**). To contextualise genome evolution across BTNR, we included a set of 74 chromosome-level genome assemblies spanning broad angiosperm diversity (“reference genomes”; **Table S3**), resulting in a comparative dataset of 573 genomes. Gene family clustering across this dataset yielded 1,339,467 orthogroups (OGs; putative gene families).

Phylogenetic reconstruction (**Fig 2; Supplementary Section 3**) using single-copy nuclear genes produced a species tree congruent with recent angiosperm-wide phylogeny based on 353 nuclear markers^22^, and provided phylogenetic placements for 52 genera previously lacking genomic resolution. Comparative time-normalised lineage through time (LTT) analyses (**Fig 2**) showed that the ancestors of the BTNR flora had increased diversification rates towards the end of the Cretaceous period, implying that their high diversity had its foundations in the angiosperm terrestrial revolution. Many BTNR families are restricted to tropics, consistent with the tropical niche conservatism hypothesis, which posits that most angiosperms originated in tropical environments and only some lineages subsequently dispersed outwards^11,13,14^.

**Figure 2:**
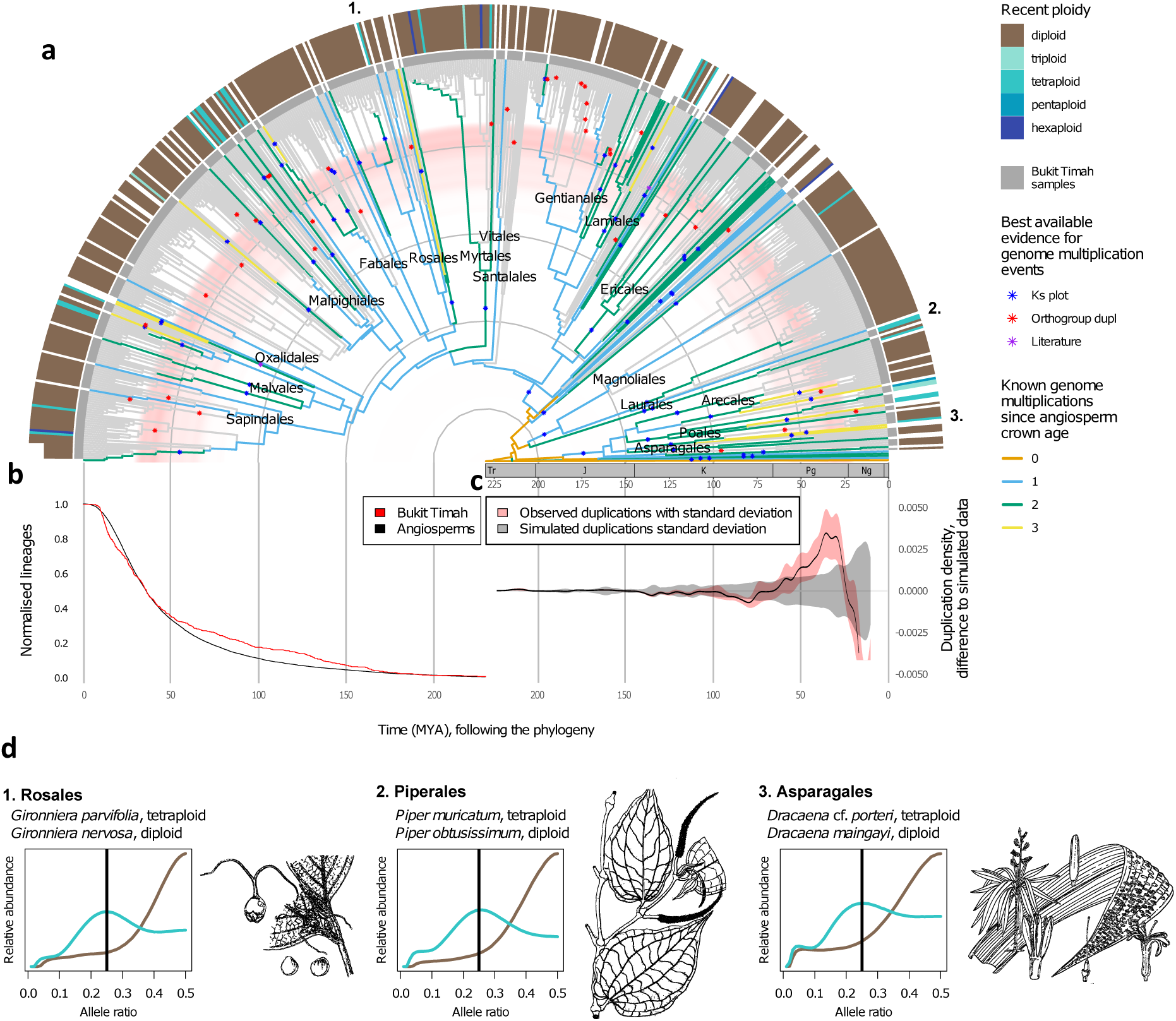
Time-calibrated nuclear gene phylogeny and genome multiplications in Bukit Timah (BTNR) angiosperms. **a)** Known genome multiplication events mapped on the phylogeny (for information on individual events, see **Table S4** and **Fig S34**. Branch colours show the entire polyploid history of those samples for which it could be tested. Neopolyploid samples are highlighted with coloured blocks. Orders with six or more species are named. Samples from BTNR are highlighted with grey blocks at the tips; the other samples are published high-quality genome assemblies that were used as references. The background colour highlights periods when gene duplications were more abundant than in simulated gene trees. **b)** Lineage through time plot showing accumulative branching of the of BTNR angiosperm phylogenetic tree, compared to a recent global angiosperm phylogeny^22^. **c)** Time periods when observed gene duplications were more abundant than in simulated uniform null model. Y-axis shows the frequency of orthogroup duplications in the observed data (standard deviation highlighted in red) subtracted from a simulated null model with uniformly distributed WGMs. **d)** Allele frequency plots of three orders with neopolyploid (cyan), partially collapsed genome assemblies, compared with a related sample that is not neopolyploid (brown). Density peak at 25% read coverage of the less-common allele on heterozygous polymorphism sites suggests the sample to be a neotetraploid with a mostly collapsed genome assembly (only tetraploid examples are shown); a peak at 50% suggests that the assembly is not collapsed and that the sample behaves as diploid, that is, it is either a diploid or allopolyploid with considerable subgenome divergence. Ng = Neogene, Pg = Paleogene, K = Cretaceous. [Drawings in public domain: Rosales: https://doi.org/10.5962/bhl.title.274; Piperales: https://doi.org/10.5962/bhl.title.911; Asparagales: https://doi.org/10.5962/bhl.title.10921]

Together, these results establish BTNR as a uniquely sampled tropical community in which both ancient diversification and contemporary coexistence can be examined in a single genomic framework. They also provide a reference baseline from which to explore how evolutionary and ecological processes have jointly shaped one of the most species-rich tropical floras documented to date.

### Limited role of polyploidy and hybridisation in tropical plant diversification

Plant genome macroevolution proceeds through two main processes, whole genome multiplications (WGMs, or, polyploidy events) followed by diploidisation^23,24^, and small-scale tandem duplications (TDs) that generate rapid local expansions of gene families^25–27^,. In the BTNR flora, gene duplications were significantly more frequent at 45.9–28.4 Mya (**Supplementary Section 4; Fig 2**), coinciding with the Eocene–Oligocene transition, a period marked by global climatic upheaval^28^ and increased migration of lineages from the Indian raft^29^. The majority of duplications in BTNR flora arose from TDs, and were enriched for gene families associated with environmental responses, particularly resistance (resistance R genes with a characteristic NB-ARC domain^30^) and key enzymes mediating secondary metabolism^31,32^; the cytochrome P450 involved in oxidation^33^ and UDP-glycosyltransferase (UGT) involved in glycosylation^34^. However, comparable temporal patterns observed in reference genomes indicate that these duplication pulses reflected global environmental shifts rather than tropical-specific processes (**Table S4)**.

Although WGMs have often been proposed as generators of angiosperm diversity, previous studies suggest that polyploidisation plays a comparatively minor role in the tropics. While estimates in temperate regions range between 20–50% of angiosperms being recent polyploids^35–37^, and that around 15% of speciation events involve a ploidy increase^38^, considerably lower rates have been indicated in the tropical forests^39^, possibly because they are dominated by perennial woody lineages where polyploidy is rare^40^.

Minor-allele-balance analysis^41–45^ revealed signatures of neopolyploidy in only 7% of species (**Fig 2)**, and even under conservative assumptions – assuming all the samples that failed quality checks (48 species) or allele balance analyses (29 species) were polyploid – the maximum possible proportion would be 20.5%. Polyploidy was rare in trees (5.6-8.6%), but substantially more frequent in herbs (46-61%), consistent with the global tendency for neopolyploidy to be associated with herbaceous life forms and stressful environments^39,40^. Importantly, polyploid species in BTNR did not possess measurable ecological or biogeographic advantages – neither in geographic range (*p*=0.85, paired two-value *t*-test) nor habitat breadth (Welch Two Sample *t*-test, *p*=0.698, **Supplementary Section 4.6**) – indicating that recent polyploidisation has not facilitated the establishment of novel niches in this community. Hybridisation, evaluated using *f3* tests^46^, was similarly infrequent, suggesting that introgression is unlikely to account for community-level diversity patterns in this forest (**Supplementary Section 5; Table S5**).

These findings provide the first community-wide estimate of neopolyploidy in a tropical forest and indicate that polyploidisation has contributed less to tropical plant diversification than to diversification at higher latitudes. In BTNR, the dominance of perennial woody species, intense competitive regimes, and limited opportunities for novel niche establishment likely constrain the persistence of polyploid lineages. As a result, the major evolutionary drivers of tropical plant diversity must lie elsewhere in the genomic architecture and selective environment of rainforest communities.

### Genome content evolves under stabilising selection of variable strength depending on gene function

To investigate the evolutionary forces shaping genome composition across BTNR, we analysed orthogroup (OG) clustering using OrthoFinder (**Supplementary Section 6**), focusing on 65,423 OGs that were present in all species among 48 well-sampled genera (≥3 species). Within this set, gene counts exhibited strong phylogenetic stratification, as on average 50% of variation was explained by phylogeny (using nested random effects model), and principal component analysis showed a grouping of species by their order (**Fig 3a**). The OGs driving these patterns were enriched for secondary metabolism, developmental processes, defence responses, symbiotic interactions and transposable element (TE) activity (**Tables S6–S7**) – categories previously associated with major morphological and ecological diversification across angiosperms^47–49^. Congruent results were obtained from both univariate analyses, identifying various developmental pathways, secondary metabolism and plant immune responses to be enriched among OGs with high phylogenetic component (**Table S8**), as well as phylogenetic ANOVA comparisons of BTNR plants against the reference genomes, with enrichments for reproductive and developmental processes, immune responses and interspecies interactions (**Table S9**). Together, this indicates diversification of these processes among BTNR plants.

**Figure 3.**
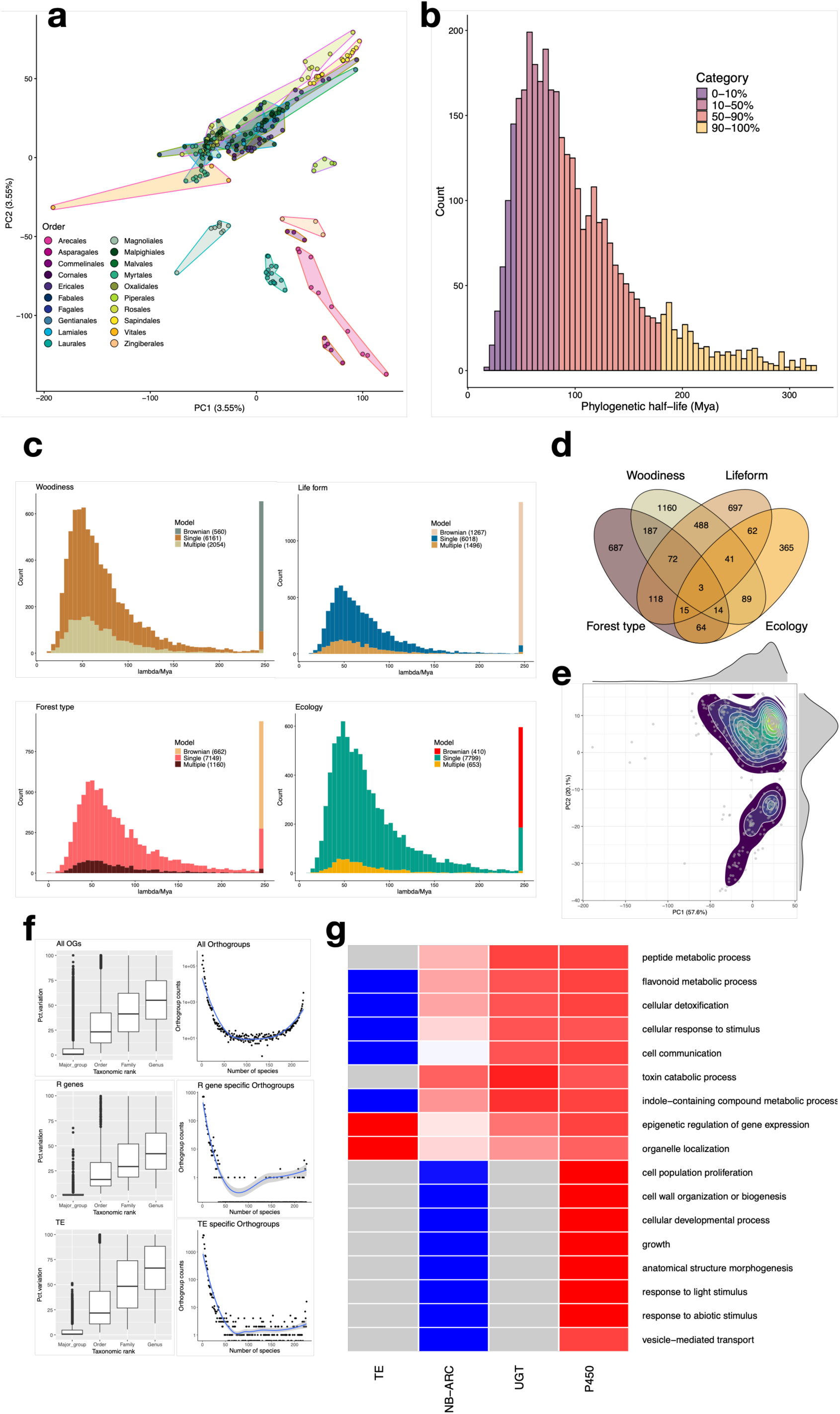
Analysis of protein-coding genes among Bukit Timah species. **a)** Principal component analysis (PCA) of the orthogroups (OGs) shared by 48 well-represented genera with at least three representative species. The coloured hulls illustrate the plant orders; the colour coding is provided in the figure legend. **b)** A histogram of phylogenetic half-lives of the OGs obtained from a univariate Ornstein-Uhlenbeck (OU) model (**Supplementary Section 8**). The different colour shadings indicate the 10, 50, 90, and 100% quantiles. The number of OGs that were better modelled as Brownian motion (in terms of AICc) are shown in the leftmost bar with a purple-coloured bar. **c)** Histograms illustrating the half-lives (lambda) from multi-mode OU modelling using four of the traits quantified in this study: woodiness, life form, forest type and ecology. The different fill colours in the histograms indicate the type of model that best explained the given OG in terms of AICc: Brownian motion, single optimum OU, or trait-specific OU. The colour palette for each of the traits is given in the figure legend, as well as the total count of each of the model categories. **d)** A Venn diagram illustrating the number of overlaps between OGs with trait-specific optima. **e)** A phylogenetic PCA visualisation of the combined set of OGs with at least one trait-specific optimum. **f)** Left: Proportion of variance explained by major group (eudicot, monocot or magnoliid), order, family, or genus. Right: Number of OGs shared with a specified number of species. Solid line illustrates loess fit, while the grey band is the standard deviation. **g)** Multivariate Ornstein-Uhlenbeck modelling identified stationary state correlations between biological processes and tandem duplicated genes (NB-ARC: resistance R genes, UGT: UDP-gelycosyltransferases, P450: Cytochrome P450) and transposable elements (TEs). For each tandem duplicate family, the top five correlations for each potential driver are shown (for a full figure see **Supplementary Section 8**). If the displayed cell is grey, the Brownian motion model better explained the data in terms of AICc.

To distinguish neutral drift-driven processes from natural selection, we modelled gene count evolution using multivariate Ornstein–Uhlenbeck (OU) processes^50–53^ (**Supplementary Section 7**), which partition changes attributable to phylogenetic drift from those driven by stabilising selection towards a shared optimum. Remarkably, 93.5% of the OGs evolved under stabilising selection (**Figure 3b**), and this pattern persisted also when genes were summarised into GO categories (97%; **Extended Data Figure 1, Supplementary Section 7.1**), implying pervasive constraints on genome expansion. These findings indicate that most functional processes are maintained near their long-term optimum states rather than drifting freely, consistent with strong selective costs associated with excessive genomic complexity.

The OU model estimates also strength of selection, described by phylogenetic half-life^53^ that is inversely proportional to the evolutionary pull towards the optimum. The half-life varied systematically across biological processes (**Figure 3b**). The top 10% of OGs experiencing the strongest pull towards the optimum (shortest phylogenetic half-lives) were involved in housekeeping processes such as highly conserved core DNA and RNA transcriptional machinery. Moderately constrained categories (10–50% quantile) were enriched for primary metabolism and cellular processes, whereas relaxed selection (50–90% quantile) was characterized by secondary metabolism, defence, development and growth-related functions, and finally the processes with the most lenient selection (remaining 10%) were linked with environment, such as environmental sensing, signaling, defence, and development (**Extended Data Figure 2)**. Again, demonstrating the robustness of our result, we obtained largely similar results when carrying out the analyses at GO level (**Supplementary Tables S10-S13**).

In sum, the model-based estimation of the strength of selection on different biological processes provides a genomic and theoretical framework for the classical observations that basic cellular machinery is deeply conserved, while environmentally responsive processes that are often implied in adaptation diversify more readily across lineages. A large body of studies on fossil data have shown morphological rates of evolution to follow a negative power law^54^, paradoxically implying that the amount of evolutionary change does not seem to depend on the time interval over which it is measured. Theoretical studies suggest that such behaviour can arise from a random walk consisting of short -term population-level adaptations tracking an optimum that depends on current environmental conditions, while over longer time scales the optimum itself would track a stable primary niche of the species^52^. Our results offer the first empirical demonstration that genome evolution in a tropical plant community is dominated by process-specific evolutionary optima, revealing that stabilising selection governs gene family evolution across ecological and phylogenetic scales. They further suggest that microevolutionary fluctuations around these optima, combined with occasional shifts in selective environments, shape macroevolutionary patterns of plant diversification, linking short-term adaptation with deep-time evolutionary outcomes, and explain why genome-level adaptation appears to slow down when inspected over longer time scales.

### Genomic convergence is driven by plant form and function

Because most orthogroups (OGs) evolve under stabilising selection, we tested whether plant traits impose their distinct evolutionary optima. We assembled trait data on life form, woodiness, forest layer (for trees), and primary versus secondary forest association. Depending on the tested trait, 7.4-23.5 % of OGs displayed distinct trait-specific optima (**Figure 3c**), and in total ∼36% of OGs showed multiple optima (**Figure 3d)**. Many of these orthogroups were highly or moderately constrained, indicating that fundamental biological pathways have converged on different genomic states in association with ecological strategies.

Phylogenetic principal component analysis of OGs with multiple optima (**Figure 3e)** illustrates that species occupy discrete regions of gene space corresponding to ecological niches—effectively forming “lumps” of convergent genomic states. Because ∼97% of the processes evolve under stabilising selection, and more than one-third of these optima are stratified by ecological traits, these findings indicate widespread convergent evolution in gene space that is driven by plant niches. Once established, these optima represent genomic attractor states that may be difficult to escape, reinforcing niche differentiation through stabilising selection on core biological processes.

Independent phylogenetic ANOVA analyses identified functional categories associated with each phenotype (FDR corrected p-value < 0.05; **Supplementary Section 7.3.6, Tables S14–S35**), and showed strong concordance with OU modelling (*p*=0.003, Fisher exact test for overlap), suggesting that trait-specific genomic optima reflect genuine evolutionary constraints rather than analytical artefacts.

Simulation models have shown that niches can emerge from neutral dynamics when resource availability fluctuates periodically, yielding “species lumps” that share similar resource-use profiles^10^. Within each niche, high similarity of species promotes coexistence, as the species are interchangeable with no serious fitness advantage over the others. Our empirical results demonstrate that such structuring also manifests at the genomic level: stabilising selection towards niche-specific attractor states partitions gene space among plant ecological strategies. Although developmental programmes appear conserved across BTNR flora, differences in phenotype are mirrored by convergent amplification of gene copy numbers^55^, indicating differential resource allocation to biological processes among ecological niches.

### Evolutionary trade-off between defence versus growth and reproduction

Since niche similarity promotes convergent evolution, we next examined processes associated with species-level divergence. Rarefaction analyses indicated that the number of new gene families (OGs) per species started to plateau but had not yet saturated (**Supplementary Section 8.1**) as, across different permutations, on average 230 species-specific OGs were still emerging in the final species added to the analysis. The results remained essentially the same when controlling for the varying taxonomic representation, as across 48 genera with ≥3 species 12.8% of all OGs were species-specific. Univariate analyses found intraspecies and host interactions as well as plant reproductive and early development processes (pollen recognition, seed germination, anatomical structure arrangement, cell cycle process) as weakly phylogenetically constrained (**Table S36**), suggesting potential roles in lineage divergence. Although most species-specific OGs lacked clear annotation, those with identifiable functions were not enriched for any single process. However, when focusing on tandem duplications and transposable-element (TE)–derived genes (**Supplementary Section 8.2;Tables S37-S38**), species-specific OGs were strongly enriched for resistance (R) genes and TEs, but depleted for P450s (Fisher exact test, all with p-value < 2.2e^-16^), indicating that R genes and TEs – but not secondary metabolism broadly – contribute to recent genomic diversification in this flora (**Fig 3f**). Furthermore, the R gene and TE OGs had generally lower than average phylogenetic contributions and the majority of them were specific to species or genera.

Phylogenetic ANOVA further identified R genes, P450s and TEs enriched among the OGs that discriminate ecological phenotypes, linking pathogen defence, developmental strategies and biochemical diversification to species-level differences (Fisher exact test, adjusted *p*-value<0.05; **Supplementary Section 8.3**). These patterns are consistent with evolutionary theory predicting trade-offs between ecological strategies – such as stress tolerance and rapid growth^56,57^ – and with empirical evidence that R genes evolve rapidly and exhibit high turnover across taxa.

To quantify these trade-offs, we modelled evolutionary correlations across all 168 top-level GO categories using multivariate OU model (**Supplementary Section 8.4**). While most relationships were positive, R genes showed strongly negative correlations with reproductive and growth processes, and positive correlations with other defence-related functions including secondary metabolism (**Fig 3; Figure S47**). These patterns indicate a pervasive evolutionary trade-off between growth and reproduction versus pathogen defence at genome scale. One interpretation is that enhanced growth and reproductive diversification reduce pathogen exposure (“escape”^58^), whereas heightened pathogen pressure favours investment in defence at the expense of developmental innovation. Defence allocation therefore appears to divert adaptive potential from growth-related pathways, potentially desynchronising evolutionary tracking of developmental optima among coexisting species.

Life-history correlates reinforce this interpretation. R gene counts positively correlate with life span and stress-tolerant strategies^59^, and negatively with growth rate^59–61^. Consistent with this, trees possessed significantly higher R gene counts and higher OU optima than other life forms (pairwise Wilcoxon tests, **Supplementary Section 8.5**), supporting the hypothesis that long-lived species invest more heavily in pathogen defence^62,63^. Among trees, understorey species exhibited reduced R gene counts relative to canopy and emergent species, yet had higher inferred evolutionary optima – suggesting that current pathogen pressure may be lower than the long-term average. The depletion of large herbivores in Southeast Asian forests^64^ may partly explain these patterns.

Overall, these results provide community-scale evidence that a genome-wide defence–growth trade-off shapes evolutionary trajectories in tropical forests. By constraining developmental diversification, pathogen pressure may promote speciation through reduced tracking of growth optima, consistent with ecological hypotheses linking escape from natural enemies to accelerated lineage diversification.

### Community composition is driven by processes associated with environmental fluctuations

To examine how the long-term genomic convergence manifests in contemporary community structure, we compared primary and secondary forest habitats within BTNR. Successional dynamics in tropical forests generate contrasting ecological strategies: secondary forests are colonised by fast-growing, light-demanding pioneers following disturbance, whereas in intact primary forest, shade-tolerant canopy and emergent trees dominate over longer intervals^65^. BTNR harbours substantial areas of secondary forest that have developed on previously cleared land^66,67^, making the forest suitable for such comparisons.

Phylogenetic ANOVA identified distinct functional categories associated with each forest type (**Supplementary Section 9.1).** Primary forest species shared higher gene counts in processes linked to regulation of DNA repair, cell wall dynamics, phloem and lateral root development, whereas secondary forest species were enriched for glycosylation, auxin signalling and metabolism (Fisher exact test, adjusted *p*<0.05; **Tables S39-S40**). At the orthogroup-level, the OGs differentiating forest types were significantly enriched also for R genes (Fisher exact test, *p*=0.045), consistent with ecological differences in defence allocation across successional gradients. OU modelling revealed 364 GO categories (1,160 OGs; **Table S41**) with forest-specific optima (**Figure 3c**), including higher optimum for lateral growth developmental processes in primary forests, and enhanced metabolic and detoxification pathways in secondary forests. As there is less organic buffer material in secondary forests, the species may be more challenged by inorganic toxic compounds in the soil.

Species-level analyses, however, do not account for community structure arising from coexistence. To identify genomic processes associated with local dominance, we mapped large trees across BTNR^18^ to 100 × 100 m grid cells (**Fig 1)** and computed community-weighted gene counts, comparing observed communities against null expectations generated by random species sampling (**Supplementary Section 9.2**). When applying the analysis to R genes, the primary forest cell counts significantly exceeded null expectations, consistent with the prevalence of long-lived, stress-tolerant lineages in these habitats (**Fig 4a**). In contrast, the secondary forests were significantly depleted compared to null, reflecting emphasis on fast life cycles. Cytochrome P450 gene families also displayed strong community-level stratification: primary forest communities showed convergent enrichment of CYP81 and CYP706 families—enzymes associated with biosynthesis of defence-related sesquiterpenes^68–71^—whereas secondary forest communities exhibited higher diversity of P450 functions linked to metabolic breadth, including triterpenoid biosynthesis^72–74^, developmental hormone regulation^75,76^ and defensive alkaloids^77,78^ (**Table S42; Fig 4b**). These patterns reflect contrasting life-history strategies: primary forest communities converge on a narrower defense-centric repertoire dominated by highly versatile enzymes^70,71,79^, whereas secondary communities diversify across broader metabolic functions suitable for rapidly changing environments. Functional interpretations of these enzymes, including roles in hormonal regulation and drought responses, further support links between gene family composition and ecological strategy.

**Figure 4:**
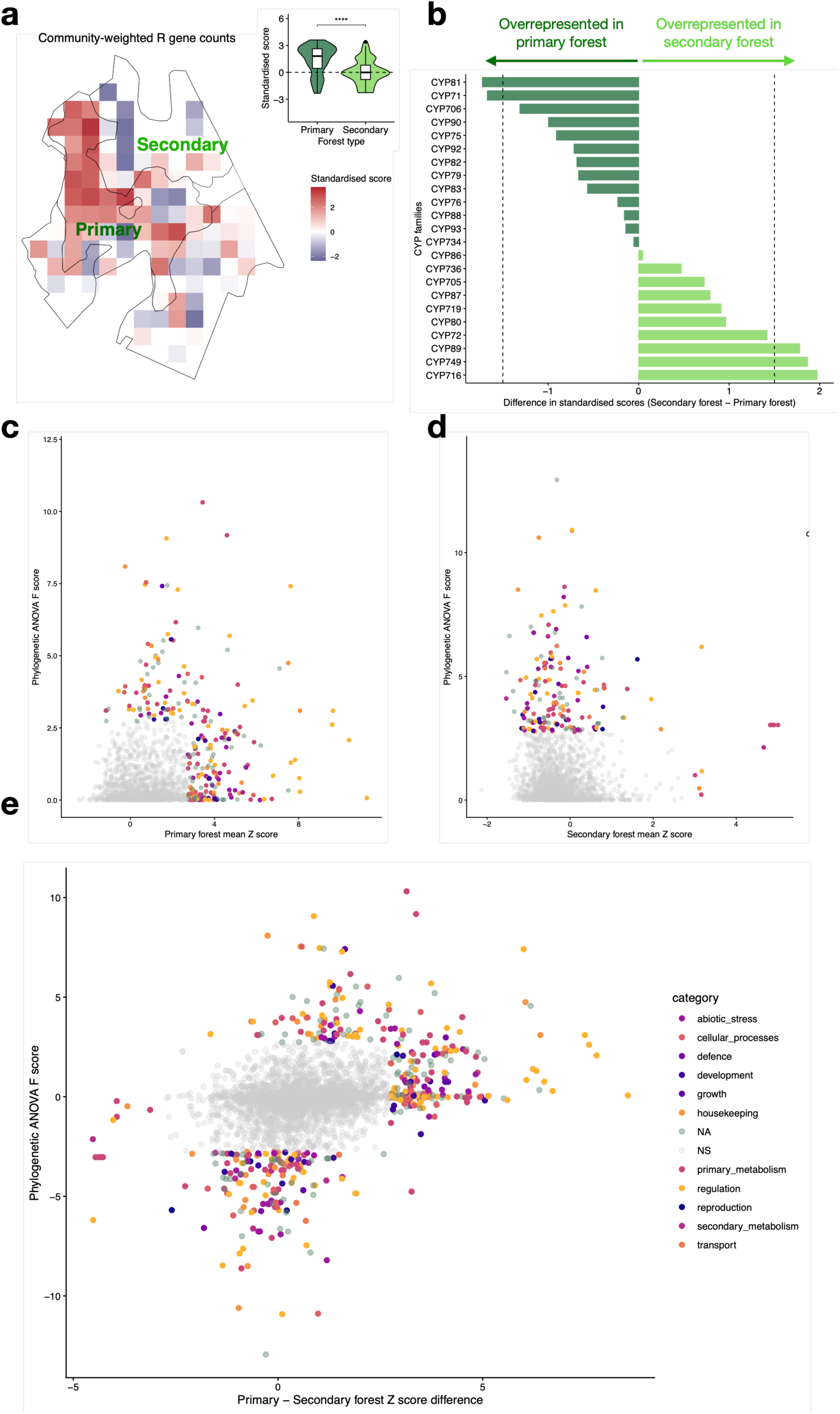
Spatial genomic analyses of R gene counts and P450 diversity with respect to stages of forest succession. **a)** Species in primary forest have greater numbers of R genes (red) than randomised community, whereas species in secondary forest have a lower number of R genes than randomised community. Violin plot of enrichment scores stratified by forest type is shown on the top right panel. **** illustrates *p*< 0.0001 according to Wilcoxon test. **b)** Bar plot of mean enrichment score differences for each P450 family by forest type. Positive scores indicate an enrichment of gene counts in secondary forest, while negative scores indicate enrichment of gene counts in the primary forest for each CYP family. Next, an analysis of community structure-driven gene set enrichment (x-axis; Primary forest mean Z score against randomised community structure) versus forest type niche-driven convergent evolution (y-axis; phylogenetic ANOVA Z score) in **c)** primary forest, **d)** secondary forest. **e)** The standardised differences between the Z scores (x-axis) versus phylogenetic ANOVA F scores on y-axis. Only the GO processes that significantly differ (p-value <0.05) from null are highlighted, the colour palette for **c**,**d**, and **e** is illustrated in panel **e.**

Extending beyond these fast-evolving gene families, for most GO categories the primary forest communities harboured more genes than secondary forests, and many of these were significantly elevated compared to a null model (**Table S43**). For most of the significantly enriched GO categories the counts were largely driven by abundant species, implying that dominant lineages disproportionately shape community genomic composition. Larger gene repertoire in the community may allow resilience under diverse environmental conditions and eventual succession over the secondary forest species. Within the communities, the strongest departures from null expectations involved regulatory processes, in primary forests these predominantly involved small GTPases and auxin signaling (**Fig 4c; Supplementary Section 9.3**) – GTPases are molecular light-regulated switches with ubiquitous roles in plant growth^80,81^ and environmental responses. As primary forest trees are shade-tolerant and can spend decades at seedling stage^82^, such molecular switches, together with auxin signaling, may facilitate shade tolerance and switching to rapid growth when light becomes available, whereas their role in drought responses may be linked with sensing of the El Niño oscillation producing drought events that regulate masting and staggered flowering in dipterocarps^83^.

By contrast, secondary forest communities were characterised by primary metabolism involved in RNA processing and translation optimisation pathways (**Fig 4d**) – processes consistent with rapid growth and flexible responses under fluctuating conditions^84^ – as well as seed longevity, in agreement with the prevalence of species forming persistent seed banks during succession^85^.

To understand how the community structure relates to processes under stabilizing selection, we compared community-level patterns with trait-driven genomic convergence. The two processes operated on distinctly non-overlapping categories (**Fig 4e**). Apparently, GO categories with genomic convergence driven by primary or secondary forest niches did not provide an advantage within a community itself. In contrast, the processes that drive community composition are linked with advantages in detecting environmental changes. Furthermore, the majority of the non-random community-level gene compositions were found to be mostly driven by abundant species, suggesting that these particular traits may facilitate their local dominance.

Together, these results show that genomic processes underpinning community assembly differ from those driving deep-time trait convergence. While trait-dependent attractor states shape genome evolution over long timescales, contemporary community composition is driven by processes associated with environmental fluctuation and enforcing local species dominance. Hence, local species dynamics appears to arise from long-term convergent evolution towards certain genomic attractor states on the one hand, and environmental fluctuations contributing towards short-term species turnover, on the other hand.

## Conclusions

Understanding why tropical forests harbour exceptional plant diversity remains a central problem in evolutionary biology. By combining community-wide genome sequencing with ecological and functional trait data from a hyperdiverse tropical forest, we show that much of the present diversity has origins in the Cretaceous radiation of angiosperms, and that contemporary diversification reflects deep evolutionary constraints acting on genome composition.

We demonstrate that genome evolution is dominated by stabilising selection towards process-specific optima, and that ecological traits divide these optima into niche-associated attractor states. These findings provide a genomic mechanism for species coexistence via niche similarity: plants occupying similar ecological strategies evolve convergent genomic compositions, being constrained by long-term selective optima imposed by their functional niches.

A genome-wide trade-off between defence and growth indicates that pathogen pressure diverts evolutionary potential away from developmental diversification, potentially promoting speciation by limiting the ability of species to track trait optima. This pattern is consistent with classical ecological hypotheses that escape from natural enemies can accelerate lineage diversification^58^ and may underlie repeated radiation events throughout angiosperm history^86,87^.

Community-level analyses reveal a second evolutionary layer, in which the genomic processes that confer competitive advantage in contemporary forests differ from those driving deep-time convergence. Primary forests show signatures of investment in functions that support sustained growth under broader environmental shifts, such as light availability, whereas secondary forests rely on pathways that enhance responsiveness to rapid fluctuations. These contrasts reflect distinct ecological strategies in the two forest types and highlight how long-term convergent evolution shapes niche structure, while short-term environmental variation favours specific genomic traits that influence present-day community assembly.

Our results establish a genomic basis for linking microevolutionary adaptation with macroevolutionary diversification across an entire tropical flora. Among various ecological theories, they best align with modern coexistence theory^9^, which posits that coexistence is maintained through stabilizing mechanisms, which promote niche differences, and equalizing mechanisms, which reduce fitness inequalities among competing species. In our system, stabilizing mechanisms arise from trait-associated genomic attractor states generated by long-term stabilising selection, which structure functional niches independently of phylogeny. Equalizing mechanisms emerge from the pervasive defence–growth trade-off. Since pathogen pressure intensifies with local abundance, widespread species necessarily allocate more of their adaptive potential to defence, limiting growth-related diversification and preventing competitive exclusion. Finally, the distinct genomic signatures observed across forest types indicate that environmental fluctuations reward different ecological strategies over time, consistent with fluctuation-dependent coexistence.

By providing the first community-scale genomic characterization of a lowland dipterocarp rainforest, this work opens a path towards a more mechanistic understanding of tropical biodiversity. Future comparisons across other biomes will test whether niche-specific genomic attractor states arise consistently across environments, or whether they are unique products of long-term evolution within humid tropical ecosystems where environmental changes are slow.

## Data availability

The sequence data is available in NCBI under a project number PRJNA917477. Raw reads are available in SRA, and the plastomes and assembled genomes with contaminant filtering are in NCBI genomes, without annotations. Annotated plastomes, original contaminant-filtered nucleotide scaffolds and their annotation files, sample-specific allele-balance and coverage plots, genus-specific f3-test figures, Orthofinder data and trait database are available in dryad (https://doi.org/10.5061/dryad.j3tx95xrf).

## Acknowledgments

The project was funded by an NParks-NTU Research Cooperative Agreement, which was co-developed on the NParks side by KBHE, DJM, and GSK and on the NTU side by VAA, CL, JS, and PRP. Additional funding was provided from the Academy of Finland (decisions 318288 and 329441 to JS), Nanyang Technological University start-up grant (to JS), MoE Tier 1 (RG82/21 to JS), NTU Presidential Post-doctoral Fellowship and NUS startup grant (to JYL), and National Parks Board Reserves Fund.

The authors thank the High Performance Computation Centre at NTU Singapore, CSC – IT Center for Science, Finland, and National Super Computing Centre (NSCC), Singapore, for computational resources, and NovogeneAIT Singapore for the sequencing services. We would like to thank SBG molecular lab staff (Khoo-Woon Mui Hwang, Felicia Tay, Neo Wei Ling and Aireen Phang) and interns and other SBG staff (particularly Parasuraman Athen and Lim Wei Hao) who helped with sample collection, identification and labwork, Serena ML Lee and Bazilah Ibrahim for help with the management of vouchers, and Soh Hwee Jin Melvin for server support at High Performance Computation Centre at NTU Singapore. We would also like to thanks Alan Schulman and Jaakko Tanskanen for help with Pfam32.0_GypsyDB.hmm.

## Author contributions

KBHE, PRP, VAA, CL, DJM, GSK and JS conceptualised the project and raised funding. DJM, GSK, VAA, CL and JS co-supervised data collection, sequencing and/or storage. MAN, LMC, PKFL, KMN, YWL, CMP collected the samples and managed the identifications. MAN, LMC, YWL, JS and JJN extracted the DNA and/or managed the sequencing orders. KMN and SKYL provided ecological monitoring data on BTNR. JYL and FWSL conducted the generation time analyses. SL provided the up-to-date names for the species. MAN, JYL, LMC, KMN, HJB, MWC, JC, RPJDK, BEEDW-D, HD, H-JE, SKG, EMG, MH, SLK, PKFL, JL-Š, SL, YWL, MSEC, HKL, LN, CMP, CP, MLR, IAS, WWS, DAS, JSS, DCT, IMT, PCvW, PW, KMW, CKY and DJM provided species-level trait information. SR conducted the genome assembly, annotation and gene clustering. MAN and JS analysed phylogenetic relationships and genome and gene multiplications. LMC and SF analysed contaminants and genomic introgression. JJN, JYL and JS conducted the spatial analyses. JJN, JJYJ and JS analysed the R gene and P450 families. JS analysed the proteome data. JS, MAN, JYL, and JJN wrote the manuscript, with input from GSK, CL, ML, VAA, PYT, CHC, TU, KBHE, DAW, SJD. The following authors prepared the supplementary materials: MAN, JYL, JJN, FWSL, MSEC, SR, JS. All authors read, commented and approved the final manuscript.

## Online Methods

### Study locality

Bukit Timah Nature Reserve (1.3550°N, 103.7783°E) is a small forest reserve situated in central Singapore (**Fig 1**). It has over 800 extant angiosperm species and contains the largest contiguous tract of primary forest in the city state^88^, plus areas of secondary forest regrowing on abandoned land formerly used for agriculture^20^. The vegetation is classified as coastal hill dipterocarp forest, a special type of mixed dipterocarp forest characterised by the abundance of the dipterocarp species, *Rubroshorea curtisii*^89^. The secondary forest areas are largely devoid of all dipterocarp species, despite decades of natural regeneration^20^. The collected samples were identified by expert botanists to mitigate the effects identified by Goodwin et al.^90^.

### Species-level data

For all species, standardised life form information and ecological traits were compiled. In addition, genus-level information on mycorrhizal associations was obtained from the FungalRoot database^91^. Local conservation statuses were obtained from the most recent National Red Data Book^92^. Tree species were additionally classified into four functional groups based on their ecology: pioneer, emergent/canopy, sub-canopy and understorey. To investigate the genomic correlates of species in different successional environments, tree species were classified into “late-successional” and “early-successional” based on their relative incidences in primary and secondary forest, respectively.

### Genome assembly

We used a minimum of 30 Gbp of 150 bp paired-end Illumina sequencing data for each sample. The genomes of the BTNR species were assembled using MaSuRCA v3.3.1^93^ with an average insert size of 350 bp and SD of 100 bp. The insert size was estimated directly from the raw reads using BBMAP v38.49^94^ before running MaSuRCA. We established automated pipelines for *de novo* assembly and annotation of the genomes by combining gene model predictions transferred from model plants with *ab initio* predictions. The assemblies were then run through several stages of quality control, including microbial contaminant removal and filtering gene predictions for transposable elements. Initial quality control suggested that 499 genomes were of relatively good quality when assessed in terms of N50 values as well as for the presence of benchmarked universally conserved single-copy genes (BUSCO^21^). The genome completeness was estimated using BUSCO v3.0.2^21^ based on the “embryophyta odb9” database. We combined this set with 74 published high-quality reference-level genomes to guide the phylogenetic analyses and to determine computationally-derived protein families, orthogroups (OGs). For PSMC analyses and for SNP-analyses, we used DUSTmasker v.1.0.0^95^ to mask low-complexity repeat regions in the entire scaffolds.

### Phylogenetic analyses

To create a phylogeny for mapping whole genome duplication events and to support other analyses, we made phylogenetic reconstructions of our genome assemblies. We combined the BTNR draft genome assemblies with high quality genome assemblies from 74 families (**Table S3**).

For nuclear phylogenies, we used the output of the BUSCO analysis. Each BUSCO entry was aligned for gene tree analysis, including maximum three paralogs per sample. The BUSCO entries were aligned as amino acid sequences using mafft v7.475^96^ and back-translated to DNA using pal2nal v.14^97^, with synonymous third base positions made ambiguous using Degen v.1.4^98^. Alignments were trimmed using trimal v1.4.rev15^99^. Each BUSCO set was analysed separately with with RAxML v.8.2.12^100^, and the resulting phylogenetic trees were summarised using ASTRAL-MP 1.1.2^101^. We also made a concatenated dataset, where a single gene copy of each BUSCO entry for each species was retained using an in-house phylogeny-aware pipeline (for each duplicated gene, the most recent common ancestor of the gene copies was determined, and the offspring gene clade with better taxon coverage was chosen for the final alignment). These single-copy alignments were concatenated using AMAS^102^ and the tree was reconstructed using RAxML v.8.2.12^100^. Time calibrated phylogeny was constructed using TreePL v.1.0^103^ with fossil calibration points from Ramirez et al.^104^. We used the fossil age as the minimum age for the clade. We assumed that the maximum age for the clades was 100 million years older. We limited the angiosperm crown age to 230–175 Mya, as this range was commonly found in a meta-analysis^105^.

### Genome duplication

The complete history of polyploidy in a genome is best inferred from the divergence spectra of synonymous substitutions (Ks) in syntenic CDS pairs (syntelogs). Self-to-self comparison of the spectra was not feasible for our dataset, as the contig lengths of most of the draft assemblies were too low to estimate synteny. We detected the WGMs from reference genomes using CoGe SynMap (https://genomevolution.org/coge/SynMap.pl) and MCScanX, sometimes with help from literature, and found out their position in our phylogeny using synteny comparisons between high-quality reference genomes and BTNR samples using MCScanX (**Supplementary Section 6.1**). We then looked for further evidence of polyploidy with methods that suited our more fragmented genomes: from normalised gene duplication rates reported by BUSCO (see above), from density of the phylogenetic position of gene duplications in OG phylogenies, and from balance of biallelic SNP coverage (allele balance); the former two methods mostly rely on fully expanded assemblies where both gene copies are annotated, and the SNP-based allele balance relies on mostly collapsed genome assemblies (e.g. recent autopolyploids). Many of the detected events had previous support from literature, and a few very ancient events that were not readily observed from the synteny plots were added based on literature alone (**Table S4**).

To detect OG duplications, we chose 1049 smallest OGs that were present in all our samples. We made phylogenetic trees using RAxML v.8.2.12 and found the node position and relative age of all ancestors of gene duplicates. This, and the simulated gene tree set to which it was compared, were done using custom pipelines (see supplementary materials). To identify whether there have been time periods when gene duplicates have been retained more than by chance, we compared the timings of gene duplication events estimated from the OG phylogenies to a null model where the gene trees were simulated with random polyploidy events, and random births and deaths of gene copies uniformly distributed in individual gene tree branches. We further estimated how much the temporal patterns are affected by tandem duplications and duplications in syntenic blocks by separating the events from reference genomes used for Ks analyses.

Allele balance analysis was done by mapping read reads to the genome assembly. We made a coverage plot of total coverage and coverage of heterozygous loci from a bam output to determine the sample-specific threshold for sequencing noise. We excluded data below this threshold, and then calculated the ratio of the less common SNP to the total coverage in heterozygous positions (examples in **Fig 2**). We only applied this method if there was a heterozygotic coverage peak that could be distinguished from noise (**Supplementary Section 6.2**). Similar methods have been used in other studies with collapsed assemblies^42–45^), but SNP-based methods appear to be less often used than alignment-free k-mer based methods (e.g. Ranallo-Benavidez et al.^106^), even though they are better suited for low coverage genomes (as single nucleotide positions have higher coverage than longer k-mers) and give additional information on assembly quality. Samples with even (1:1) ratio of reads supporting minor:major allele were considered diploid (i.e. not neopolyploid), those with mostly 1:2 ratio neotriploid, and 1:3 ratio neotetraploid etc.

### Gene model prediction and functional annotation

Gene models were predicted based on a custom annotation pipeline. Homology based gene prediction was carried out using GeMoMa v1.6.1 based on the reference gene models of two model species, *Arabidopsis thaliana* and *Populus trichocarpa*. *Ab initio* gene prediction was carried out using Augustus v3.3.2^107^ trained on the BUSCO genes of the respective species and the self-training GeneMark-ES^108^. All these predictions were then combined into a single high confidence gene prediction set using Evidence Modeler v1.1.1^109^.

The predicted genes were blasted against *Arabidopsis thaliana* genes models using Blastp v2.13.0+. Using an in-house R script, the best hits were extracted with an e-value cutoff of 1e-05 and finally mapped to the predicted gene models. The corresponding model GO categories were also mapped to the gene models.

### Repeat removal

The annotations obtained from Evidence Modeler were screened for repeat elements using hmmsearch in HMMER V3.2.1 with hmm profiles collection Pfam32.0_GypsyDB.hmm from REPET database^110^. Genes that matched to repeat domains were then filtered to create a repeat-free gene set.

### Gene family clustering

The predicted proteomes were clustered together with the proteomes from the reference species using Orthofinder v2.3.3^111^ with default inflation parameter (1.5).

### Phylogenetic models

The impact of tree trait on gene counts was tested using linear mixed models with nested random effects for order, family and genus. The model was run separately for each OG; since the dependent variable was counts, we took a square root of the counts to correct for the heteroscedasticity in the data. The OGs where the marginal variance explained was greater than 5% were selected for further inspection. Tukey honest significant differences contrasts were run for each OG and the significant comparisons (*p*<0.05) were selected.

### Gene ontology enrichment

To obtain functional interpretation, the genes in all significant OGs in each comparison were collected and gene ontology enrichment analysis was run using Goatools^112^.

### Multivariate Ornstein-Uhlenbeck modelling

R library mvMORPH was used of fitting the Ornstein - Uhlenbeck (OU) model. The OU process is a stochastic mean reverting-process with three parameters, the rate of reversion to the mean, alpha, the mean value of the process, mu and the variance of phylogenetic Brownian motion model, sigma. For each GO category and gene set, both OU and a Brownian motion models were fitted, and the model with the lower Akaike AICc score was selected as the model that better explained the data. Phylogenetic half-lives were calculated for the categories where OU model yielded lower AICc value using formula H=log(2)/alpha, where H is the half-time of the mean-reversion, phylogenetic half-life. For testing different evolutionary optima for the different categorical traits, the trait was mapped to phylogenetic tree, and OU models for multiple optima, single optimum, as well as Brownian motion model were fitted to the data, the model with the lowest AICc value was selected as the best explaining model.

Correlations between traits were studied with multivariate OU model by calculating pairwise correlations with level 1 and 2 GO categories as well as the potential driver genes (R genes, cytochrome P450s, UDP glycosyltransferases and transposable elements). Model selection was done by choosing the model with the lowest AICc score. Stationary correlations between the traits were then obtained for the OU models.

