## Extended data figure for "Stabilising selection and ecological trade-offs underpin coexistence in a tropical flora"

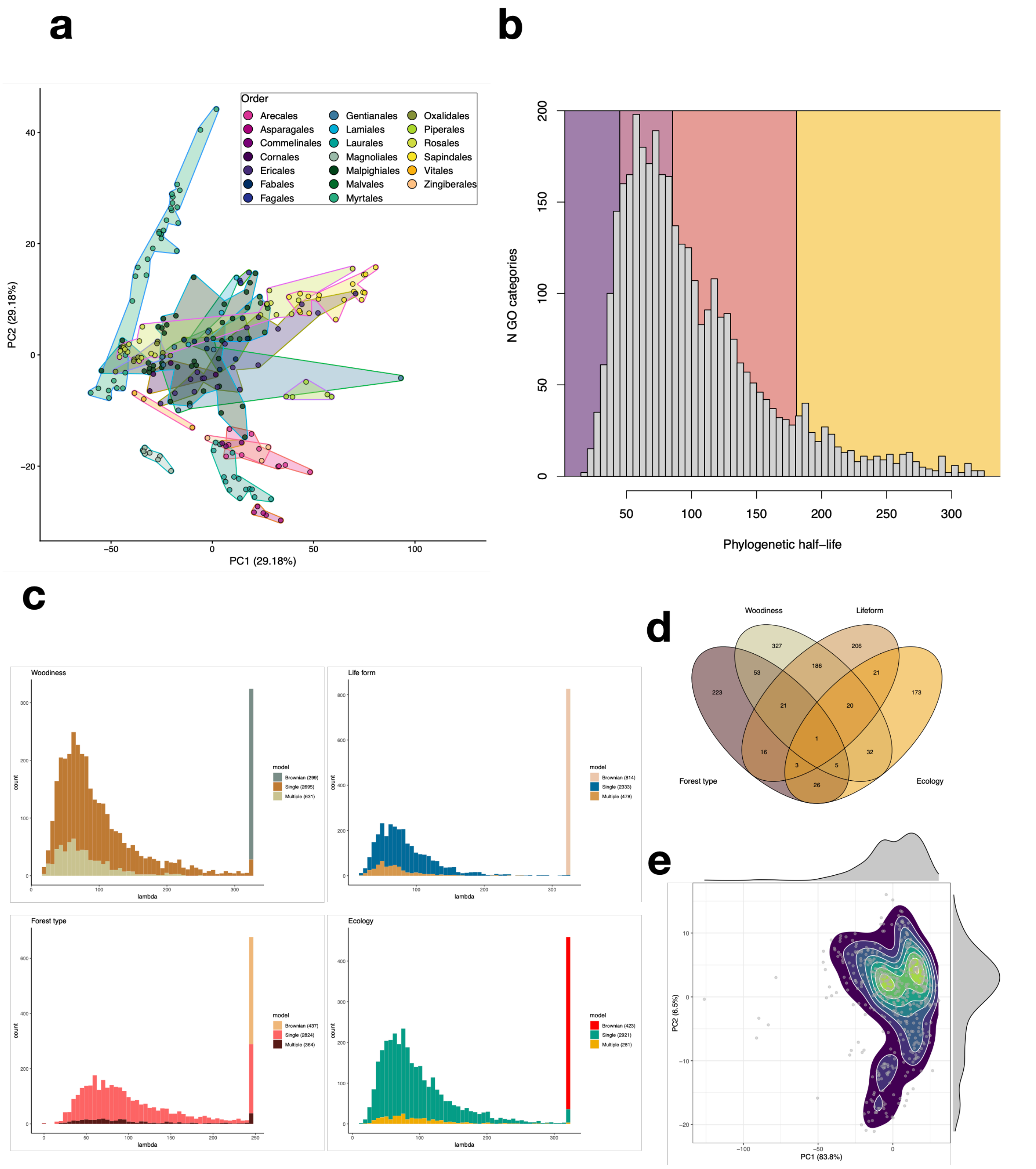


**Extended Data Figure 1. Analysis of protein-coding genes among Bukit Timah species when summarized to Gene Ontology categories. a)** Principal component analysis (PCA) of the Gene ontology categories (GOs) present in >90% of the species. The coloured hulls illustrate the plant orders; the colour coding is provided in the figure legend. **b)** A histogram of phylogenetic half-lives of the GOs obtained from a univariate Ornstein-Uhlenbeck (OU) model (**Supplementary Section 8**). The different colour shadings indicate the 10, 50, 90, and 100% quantiles. The number of GOs that were better modelled as Brownian motion (in terms of AICc) are shown in the leftmost bar with a purple-coloured bar. **c)** Histograms illustrating the half-lives (lambda) from multi-mode OU modelling using four of the traits quantified in this study: woodiness, life form, forest type and ecology. The different fill colours in the histograms indicate the type of model that best explained the given GO in terms of AICc: Brownian motion, single optimum OU, or trait-specific OU. The colour palette for each of the traits is given in the figure legend, as well as the total count of each of the model categories. **d)** A Venn diagram illustrating the number of overlaps between GOs with trait-specific optima. **e)** A phylogenetic PCA visualisation of the combined set of GOs with at least one trait-specific optimum.

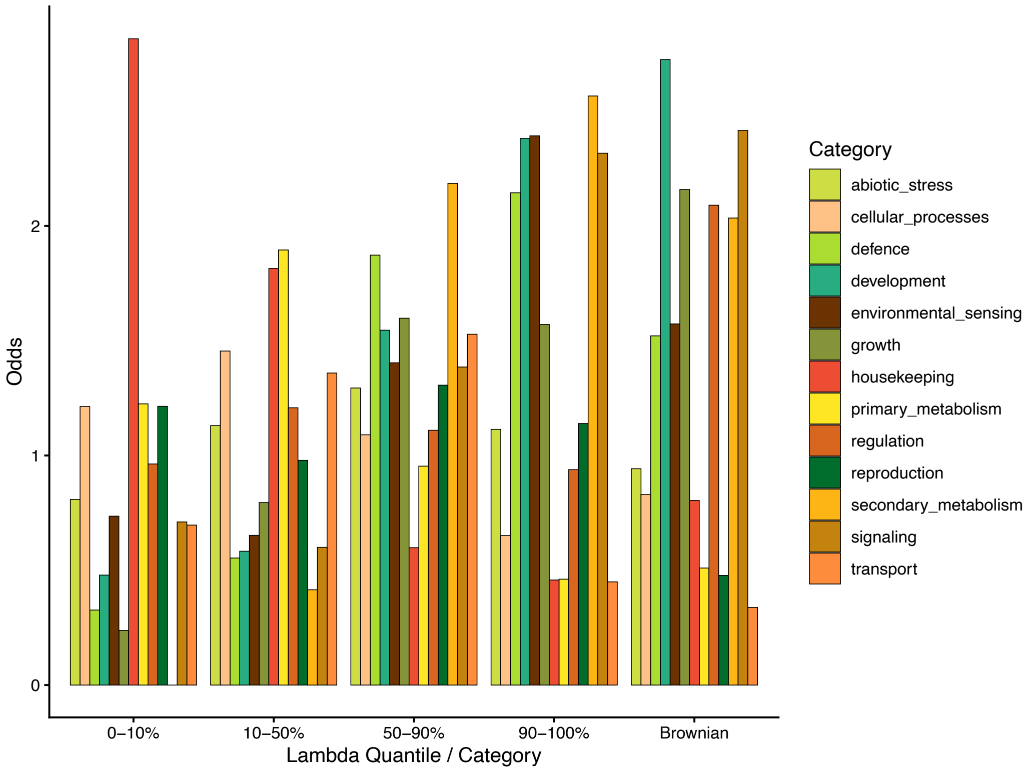


**REPRODUCTION**: reproduction, reproductive, sexual reproduction, meiosis, meiotic, meiotic cell cycle, synaptonemal, chiasma, recombination, chromosome segregation, sister chromatid, gamete, gametophyte, sporogenesis, microsporogenesis, megasporogenesis, pollen, pollen sperm, anther, tapetum, pollen tube, ovule, fertilization, double fertilization, zygote, endosperm, seed dormancy, seed maturation, seed development, embryo development, gametogenesis

**DEVELOPMENT**: development, developmental, morphogenesis, organogenesis, histogenesis, pattern formation, axis specification, cell fate, specification, identity, meristem specification, flower, floral, inflorescence, organ development, tissue development, fruit development, ripening, abscission, senescence, cotyledon, seed coat, integument, nectary, secondary cell wall, cell wall biogenesis, cell wall organization, differentiation, formation, patterning, shaping, aging, ageing, maturation, acquisition of seed longevity, seed germination

**GROWTH**: growth, cell expansion, elongation, meristem growth, root growth, shoot growth, inflorescence meristem growth, floral meristem growth, pollen tube growth, seed growth, fruit growth, cell enlargement, cell size, cell wall loosening, biomass accumulation, shade avoidance, regeneration

**DEFENCE**: defense, defence, immune, immunity, innate immune, immune response, hypersensitive response, systemic acquired resistance, induced systemic resistance, pattern recognition, pamp, antiviral, rnai, rna interference, programmed cell death induced, pathogen, bacterium, fungus, oomycete, virus, insect, nematode, herbivore, xenobiotic, toxin, detoxification

**ABIOTIC_STRESS**: response to abiotic, stress, heat, cold, freezing, drought, desiccation, water deprivation, salt, salinity, osmotic, flooding, hypoxia, anoxia, oxidative, reactive oxygen species, ros, singlet oxygen, photooxidative, high light, low light, photoinhibition, uv, radiation, ozone, carbon dioxide, ph, metal, cadmium, copper, zinc, arsenic, aluminum, starvation, nutrient deprivation

**PRIMARY_METABOLISM**: catabolic process, primary metabolic process, carbohydrate, sugar, polysaccharide, cellulose, hemicellulose, xylan, pectin, glycolysis, gluconeogenesis, tricarboxylic acid, tca, respiration, electron transport, energy generation, photosynthesis, photorespiration, amino acid metabolic, amino acid biosynthetic, lipid metabolic, fatty acid, nucleotide metabolic, purine, pyrimidine, cofactor metabolic, vitamin metabolic, glutathione, redox, starch, TCA, citrate, malate, photosynth, photoresp, pentose−phosphate, glycolytic, glyoxylate, 2−oxoglutarate, Alanine, Arginine, Asparagine, Aspartic acid, Cysteine, Glutamine, Glutamic acid, Glycine, Histidine, Isoleucine, Leucine, Lysine, Methionine, Phenylalanine, Proline, Serine, Threonine, Tryptophan, Tyrosine, Valine, amino acid, fatty, lipid, sterol, glycerol, acetyl−CoA, nucleotide, nucleoside, ATP, ADP, AMP, GMP, IMP, UMP, TMP, TDP, UDP, UTP, CTP, GTP biosynthetic, GTP catabolic, NAD, NADP, FAD, ubiquinone, coenzyme, porphyrin, heme, sucrose, trehalose, monosaccharide, fructose, glucose, galactose, mannose, maltose, glycogen, xylulose, fucose, inositol, glycosaminoglycan, glucosamine, N−acetylglucosamine, glucuronate, glucan, urea, argininosuccinate, sulfate, alcohol, ethanol, aldehyde, acetate, energy, precursor metabolites, salvage

**SECONDARY_METABOLISM**: secondary metabolic process, specialized metabolic process, phenylpropanoid, flavonoid, anthocyanin, chalcone, lignin, lignan, coumarin, terpenoid, isoprenoid, monoterpen, sesquiterpen, diterpen, triterpen, glucosinolate, alkaloid, polyketide, phytoalexin, camalexin, carotenoid, xanthophyll, wax, cutin, suberin, sporopollenin, green leaf volatile

**CELLULAR_PROCESSES**: cell cycle, mitotic, meiotic, cytokinesis, cell division, chromosome organization, chromatin, centromere, telomere, spindle, microtubule, cytoskeleton, organelle organization, membrane organization, programmed cell death, GTPase, small GTPase, Ras, Rab, Rho, Ran, Arf, dynamin, autophagy, mitophagy

**TRANSPORT**: transport, transmembrane transport, vesicle−mediated transport, endocytosis, exocytosis, secretion, export, import, recycling, homeostasis

**HOUSEKEEPING**: dna replication, dna repair, dna recombination, transcription, rna processing, rna splicing, DNA, RNA, rrna processing, trna processing, translation, repair, protein modification, protein phosphorylation, ubiquitin, proteolysis, protein catabolic process, recombinational, chromosome maintenance, telomere maintenance, chromatid, epigenetic, methylation, acetylation, pseudouridine, depalmitoylation, intron splicing

**REGULATION**: regulation of, positive regulation, negative regulation

**SIGNALING**: signaling, signal transduction, receptor signaling pathway, checkpoint signaling, kinase, phosphorylation, auxin, indoleacetic acid, cytokinin, gibberellin, abscisic acid, ethylene, jasmonic, salicylic, brassinosteroid, strigolactone, communication, MAPK

**ENVIRONMENTAL_SENSING**: response to, acclimation, adaptation, tolerance, detection, sensing, perception, tropism, phototropism, gravitropism, photoperiodism

**Extended Data Figure 2. Enrichment of plant functional categories among GO terms grouped according to their phylogenetic half-life, quantiles: <10%, 10-50%, 50-90%, 90-100%.** The bar plot shows Odds values from Fisher exact test for each of the functional category, with "1" corresponding to null model. Functional categories were obtained prior to the analysis by assigning Gene Ontology terms into 13 main categories according to keywords in the GO term name.
